# The TissueTractor, a device for applying large strains to tissues and cells for simultaneous high-resolution live cell microscopy

**DOI:** 10.1101/2024.06.28.600827

**Authors:** Jing Yang, Emily Hearty, Yingli Wang, Deepthi S. Vijayraghavan, Timothy Walter, Sommer Anjum, Carsten Stuckenholz, Ya-Wen Cheng, Sahana Balasubramanian, Adam V. Kwiatkowski, Lance A. Davidson

## Abstract

Mechanical strain substantially influences tissue shape and function in various contexts, from embryonic development to disease progression. Disruptions in these processes can result in congenital abnormalities and short-circuit mechanotransduction pathways. Manipulating strain in live tissues is crucial for understanding its impact on cellular and subcellular activities. Existing tools, such as optogenetic modulation of strain, are limited to small strain over limited distance and durations. Here, we introduce a high-strain stretcher system, the TissueTractor, designed for high-resolution spatiotemporal imaging of live tissues, enabling strain application varying from 0% to over 150%. This system is needed to unravel the intricate connections between mechanical forces and developmental processes. We demonstrated the stretcher with *Xenopus laevis* organotypic explants, human umbilical endothelial cells, and mouse neonatal cardiomyocytes to highlight the stretcher’s adaptability. These demonstrations underscore the potential of this stretcher to deepen our understanding of the mechanical cues governing tissue dynamics and morphogenesis.

**Graphical Abstract:** 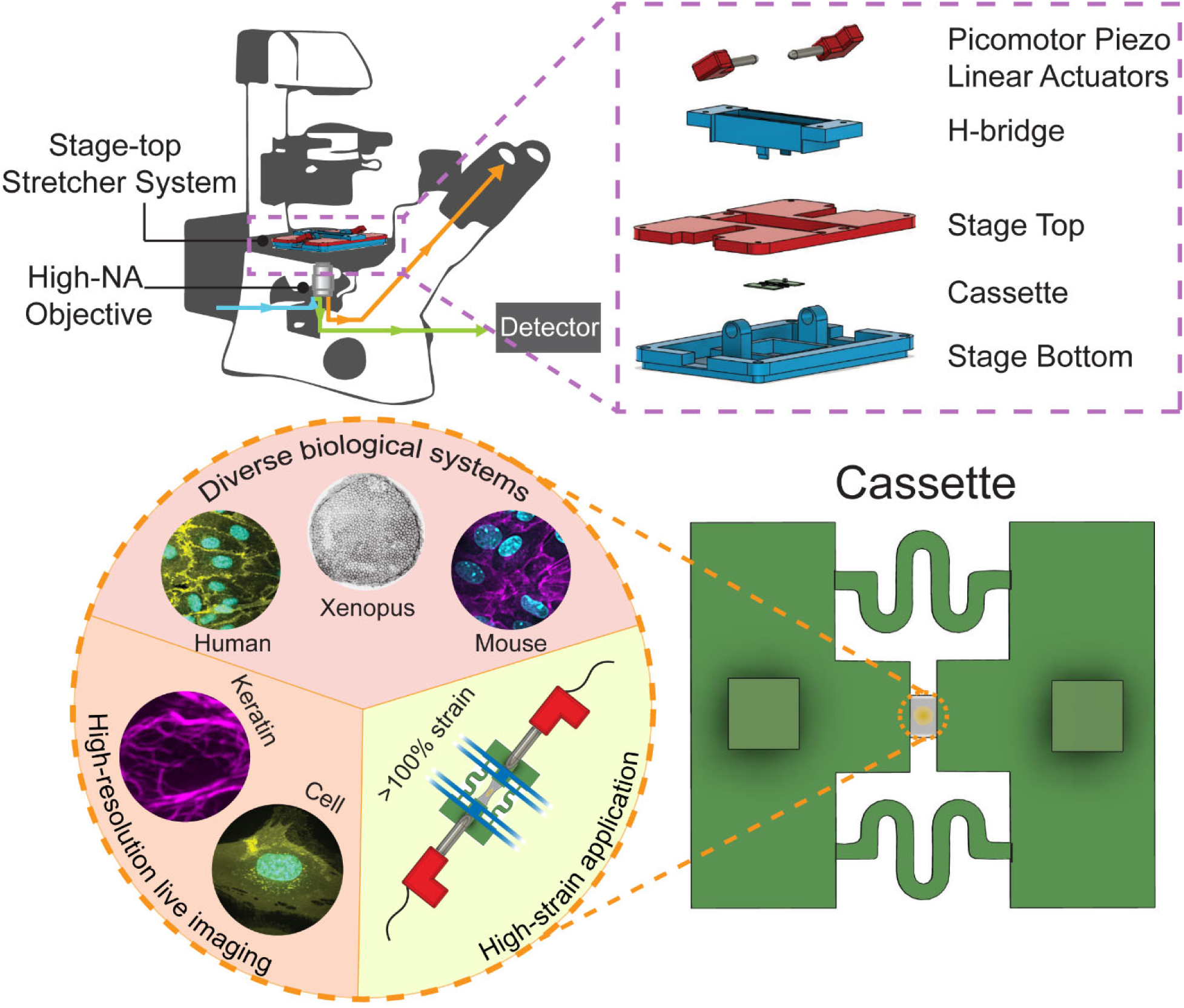

## 1. Introduction

Physical mechanics plays a critical role in orchestrating tissue shape and subsequent function across a range of contexts including early embryonic development and later organogenesis, carcinogenesis, wound healing, and regeneration.^1–12^ Mechanical strain, induced both by the local microenvironment and external sources, can guide biological processes such as cell migration, proliferation, cell fate change, etc.^8,9,13–17^ For instance, externally-applied compressive forces can reduce cancer cell proliferation and induce apoptosis,^11,18^ and in other cases, compressive forces can guide growth cone migration and drive collective neural crest cell migration.^17,19^

Lesions to mechanical processes may lead directly to birth defects via inappropriate strain and tissue malformation or by altering normal mechanically-triggered biological processes. These lesions can drive structural birth defects in organ formation, such as congenital heart defects, spina bifida, and ventral body wall closure.^8,17,20^ For instance, changes in embryo bulk mechanical properties can facilitate or delay the mesenchymal-to-epithelial transitions in *Xenopus* heart progenitor cells. These mechanical changes can cause cardiac defects; specifically, increased external mechanical tension is shown to induce 50% more mesenchymal-to-epithelial transition in heart progenitor cells, giving rises to cases of cardiac edema.^8^ Another study found that lack of tissue strain during gastrulation disrupted planar-cell-polarity in the ciliated epithelium of *Xenopus* embryos; importantly, this defect was rescued by applying exogenous strain similar to the normal gastrulation strain.^21^ Understanding how mechanics in general, and mechanical strain in particular, function in developmental processes is key to identifying underlying causes of diseases and birth defects.

High strain can also change tissue mechanical properties by fluidizing or solidifying tissues, facilitating transitions between so-called solid-like and fluid-like states by “unjamming” or “jamming” cells in the tissue.^22–25^ Such transitions may involve alterations in adhesive junctional complexes between cells to allow remodeling.^26^ Recent studies have described “jamming” and “unjamming” tissue behaviors during embryonic development,^22,27^ but it remains unknown whether or how mechanical strain alters cell-cell junctions enabling transitions between fluid- and solid-like states. Nonetheless, the capacity of a tissue to remodel is critical for its ability to dissipate strain energy, and the mechanical cues those strains encode.

Establishing causal relationships between mechanical cues and their effects on multicellular tissues requires tools capable of experimentally generating temporally and spatially defined strains, *e.g. externally controlled strain rates*, that are compatible with high resolution live-cell imaging. Key cellular and extracellular features including the nucleus, cytoskeleton, cell adhesions, and extracellular matrix have all been implicated in establishing mechanical properties as well as in transducing strain cues into signal transduction pathways. High resolution live-cell microscopy combined with image analysis pipelines can quantify the distribution of polarity factors; the dynamics of the cytoskeleton, adhesion, and membrane remodeling; and how those processes are coupled to signal transduction. The ability to control tissue strain over minutes to hours akin to controlling gene activity has immediate applications in studying the influence of mechanical cues in remodeling both synthetic and native tissues within complex micromechanical microenvironments.^28–32^

In order to test the physiological roles of strain, a tissue stretcher must be able to apply the large strains observed during embryonic morphogenesis to tissue samples cultured *ex vivo*. During the most rapid phases of embryonic morphogenesis, tissues experience large strain, ranging from 50% to greater than 100%.^21,33^ For example, tissues undergoing convergent extension exhibit more than 2-fold changes in length, greater than 100% strain, during zebrafish and *Xenopus* gastrulation and neurulation.^1,33–36^ To replicate these high *in vivo* levels of strain, we sought a stretcher system capable of reaching 100% strain or more. Furthermore, simultaneous observation of intracellular cytoskeletal and adhesion dynamics requires high numerical aperture oil immersion objective lenses that typically have small working distances that require tissues to be less than 200 µm from the coverslip. To acquire high-resolution imaging sequences while applying strain, stable tissue mounts must minimize out-of-plane torsion that would otherwise drive samples out of plane beyond the objective’s working distance. Finally, the total mass of the stretcher device must be compatible with piezo or galvo-driven z stages that are commonly used in rapid confocal sectioning in live cell imaging systems (e.g. Leica, Zeiss, Nikon, and Thorlabs).

Previously developed stretcher systems have been used to apply strains to living tissues in combination with microscopic analysis.^37^ Simple stretchers for suspended cell monolayers consist of wire cantilevers.^38,39^ More sophisticated uniaxial or biaxial stretchers use clamps or posts to bond tissues to motorized actuators.^40–42^ Clamp- and actuator-based systems are typically bulky and weigh considerably more than the 250 g mass limit of fast z-scanning stages used for high resolution confocal sectioning. Another technique to apply strain is to induce compression along one axis and thus generate tensile strain along the other two axes.^43,44^ Although this technique is easy to implement, the technique typically achieves only small strains and is limited to larger bulk tissue samples such as whole embryos. Indentation of an elastic substrate with seeded cells or tissues is also commonly used to induce tensile strain. In these indentation devices, an elastic substrate is fixed on posts, and an indenter is used to press on the substrate to deform the substrate, generating strain on seeded cells.^45^ Indentation requires steric access for positioning and travel along the z-axis, which can limit access for high resolution optics.

Past stretchers have also been designed to generate strain on cells or tissues by bonding or attaching them to an elastic substrate. The earliest efforts to stretch embryonic tissues used rubber substrates.^46^ Commercial systems (e.g., CellScale^47^) use posts to fix the edges of an elastic substrate, with strain subsequently applied by a linear actuator along the edge of the device. Another commercial device (e.g., FLEXCEL^48^), uses a macro-scale indenter to induce strains up to 30%. These systems described above have advantages and limitations, but none are well suited for high resolution confocal live-cell microscopy.

To overcome the limitations of previous stretcher designs, we developed a stretcher, the TissueTractor, capable of inducing high strain on living tissues that would also allow imaging at high-resolution on an inverted confocal microscope. Our design has three main components: an easily interchangeable cassette, motorized actuators, and a customed microscope stage insert. The modular design of our stretching system enables integration with an inverted compound microscope equipped for high-resolution confocal imaging. The cassette-based design allows simple exchange of samples for technical and biological replicates. Furthermore, the unique cassette design allows us to image samples directly through a simple cover glass, instead of support substrates, such as an elastic substrate, which are not optimized for high-resolution imaging. Additionally, the cassette design is easily modified to accommodate diverse experimental models. In this study, we first demonstrated use of the stretcher device with organotypic explants from *Xenopus laevis* embryos. Stretching tissues with the device allowed us to quantify cell strain heterogeneity across the tissue and to visualize remodeling of intracellular keratin filaments under tension. Next, we demonstrated the broader applicability of the system by modifying cassettes for use with human umbilical endothelial cells (HUVECs) and mouse neonatal cardiomyocytes, permitting observations of cell morphological changes under large strain. In conclusion, our innovative stretcher system enables high-resolution confocal imaging of living tissues under high strain from diverse animal model. This flexible and customizable system is a powerful tool for gaining insight into how mechanical cues function in remodeling tissues.

## 2. Results

### 2.1. Development of a customized stretcher system for high-resolution live imaging

To investigate how multicellular tissues respond to applied strain, we developed a microscope stage-top stretcher integrated with commercial inverted brightfield and confocal microscopes to generate large effective mechanical strain in live samples (**Figure 1a**). The stretcher system features three main components: a cassette holding live samples, motorized linear actuators to deform the cassette, and a microscope stage insert to integrate actuators with the cassette (**Figure 1b-d**). The cassette provides stability along the z-axis during stretching while allowing bilateral elastic deformation along the stretch axes (the x-axis). The cassette was fabricated out of a rigid thin polyester (PES) sheet using a 2D cutter. The cassette efficiently transmits tension to tissues attached to an elastomeric material, poly(dimethysiloxane) (PDMS), that spans a gap in the center of the cassette (**Figure 1b**, **and 1c**). Since tissues and cell thickness can vary, we match the sample thickness with the thickness of the PES sheet used to build the cassette. We find PES sheets from 38 to 127 µm thick can accommodate the thickness of samples so they remain within the working distance of a high numerical aperture objective lens. The stiff PES sheet provides a rigid mount for the two ends of the PDMS substrate. Samples attached to substrate are physically coupled to the cassette; thus, as the cassette is stretched, samples are deformed. Our design uses 2D springs cut into the PES cassette. Spring designs were selected to minimize out-of-plane motion^49^ and to stabilize the tissue sample in the field of view during a stretch cycle. Cassette rigidity and consistent bonding of the PDMS substrate to the cassette was achieved using a “sandwich” design, with a dumbbell shaped PDMS substrate bonded between the top and bottom PES sheets using UV-curable optical adhesive (**Figure 1c**). The dumbbell shape of the PDMS limited bond rupture and slippage from the PES at large strain (**Figure 1c’**). Lastly, two 3D-printed photopolymer resin block abutments mounted to the cassette allow direct contact between the two ends of the cassette and stage top apparatus (**Figure 1c**). A fully assembled cassette has a 1.8 mm wide gap, e.g. grip-to-grip (**Figure 1c**).

**Figure 1.**
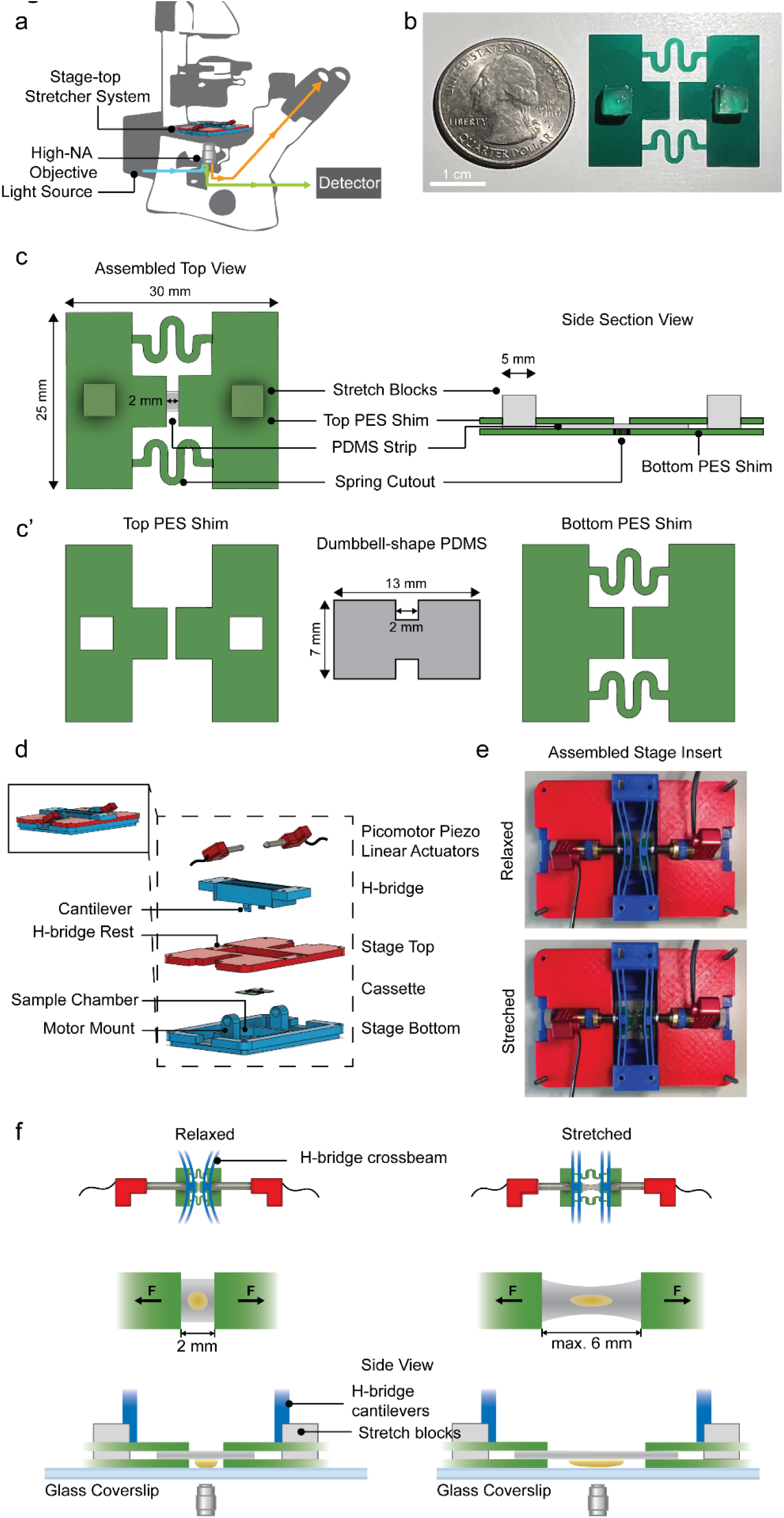
Structural configurations of the stretcher system. (**a**) a schematic of the stage-top stretcher system on an inverted microscope. (**b**) a disposable cassette. Scale bar = 1 cm. (**c**) top and side section views of the assembled cassette. The cassette is 30mm width x 25 mm length with 2 cubic stretcher blocks. (**c’**) The cassette has a top PES shim with two square cut-outs for stretcher blocks, a dumbbell-shape PDMS substrate with a 1.8 mm exposed region after assembly, and a bottom PES shim to provide xyz-stability. (**d**) an exploded view of all components in the stretcher system. From top to bottom: two picomotor piezo linear actuators, an “H-bridge” with extended cantilevers is controlled by the movement of actuators, a stage top that has a H-bridge rest to ensure a tight fit, a disposable cassette, and a stage bottom that has two motor mounts to mount the linear actuators, and a stage chamber that can be filled with liquid. All components except actuators and cassette are 3D-printed using Fused Deposition Modeling (FDM). (**e**) Top view of the assembled stage insert at a relaxed state (top) with H-bridge crossbeams bending inward, and at a stretched state (bottom) with H-bridge crossbeams straighten. (**f**) a schematic of the cassette at a relaxed state with linear actuators pushing in to bend the H-bridge crossbeams (top left); as the arms of the actuators retracted, H-bridge crossbeams become straight and stretch the cassette (top right). A schematic of a tissue sample attached to the PDMS substrate of the cassette at a relaxed state (middle left) and at a maximum stretched state (middle right). A side view of a cassette being stretched and imaged on an inverted scope, with the tissue sample faces towards the objective lens (bottom). The H-bridge cantilevers snap onto the stretcher blocks and push outward as the actuator arms retract.

The custom, 3D-printed microscope stage insert for the stretcher and cassette was designed to fit commercial inverted brightfield or confocal microscopes (**Figure 1a**, **and Figure S1**). To achieve an optimal combination of high-resolution live imaging and effective mechanical manipulation of biological samples, the stage insert has four components: a stage insert bottom (specific to the microscope), a stage top, an “H-bridge” cantilever, and two Piezo linear actuators (**Figure 1d**). The bottom includes a 45 x 50 mm sample chamber and two motor mounts for securely mounting motorized linear actuators, provides stability and prevents shifting during live imaging. The H-bridge consists of two cantilevers with extensions to contact the two abutments on either side of the cassette (**Figure 1d**, **and 1f**). The top of the H-bridge aligns with the two linear actuators so each motor is able to independently displace one cantilever (**Figure 1f**).

Stretch occurs as the crossbeams of the H-bridge are displaced, transmitting displacement to the two abutments on either side of the attached cassette. In preparation for uniaxial stretching, the two linear actuators are extended to deflect the crossbeams of the H-bridge inward. Next, the cassette is placed in a relaxed state into the culture-media filled sample chamber, with the coverslip serving as the bottom of the chamber, allowing direct and unimpeded microscopy of the sample (**Figure 1f**). The abutments of the cassette are aligned with, but not yet contacting, the lower extensions of the H-bridges. At this point, the relaxed cassette is connected to H-bridges by retracting the actuators. As the crossbeams of the H-bridge relax, they contact the cassette abutments (**Figure 1e-f**, **and Supplementary Video 1**). A small initial uniaxial bilateral stretch of the cassette immobilizes the sample within the field of view (**Figure 1f**). Motors, H-bridge cantilevers, and cassette with a PDMS substrate are rigidly integrated. Large scale strain is applied in steps as the cassette ends are moved apart and the PDMS stretches. The exposed PDMS substrate within the cassette can be stretched to a maximum of 6 mm, generating up to 200% strain (**Figure 1f**). Cassette end movements are controlled either by computer-controlled positioning of the linear actuators (LabView), or manually via a joystick. Image acquisition is carried out with microscope automation software (µManager 2.0^50^).

Since large displacements can generate apparent drift of the image across the field of view, we developed a custom microscope control AutoCenter to recenter the image. AutoCenter was based on the built-in autofocus plugin of our microscope automation software (µManager 2.0^50^). In brief, AutoCenter runs between actuator movements and adjusts the position of the XY stage to counter drift of the sample. The plugin module is executed at the beginning of each specified acquisition timepoint with two user inputs: (1) the channel to use for the adjustment, and (2) a search range along the Z-axis (usually equal to the Z-stack range set at the beginning of the acquisition). Using the user-input search range, AutoCenter will find the Z position that exhibits the greatest sharpness and capture a reference image. The module will then calculate the xy displacement between the new reference image (from the current time point) and the previous reference image (from the last time point) using conjugate multiplication of the fourier transforms of the two images, inverse transforming the result, and then finding the Δx, Δy deviation of the brightest pixel from the center of the image. Δx and Δy are used to set the stage to the new position in register with the earlier time point (**Figure S4, and Supplementary Video 3**).

### 2.2. The strain profile of the stretcher system

Substrate strain uniformity is critical to uniform deformation across attached tissues. To validate strain uniformity, we coated PDMS in the cassette with 5 µm-diameter green fluorescing polymer beads (**Figure 2a**). The linear actuators were controlled by the LabView program to stretch the cassette at a constant velocity of 52 ± 3.5 µm per minute for 45 minutes (**Figure S2**) and the cassette was imaged every 30 seconds (**Supplementary Video 2**). To verify the strain profile across the substrate, we selected two locations: (1) near the site where the PDMS substrate connects with the PES sheet and (2) at the center of the PDMS where we intend to track tissue strain (**Figure 2a**). We tracked the fluorescent beads every 15 minutes during stretch to investigate the strain uniformity across the PDMS substrate (**Figure 2b**).

**Figure 2.**
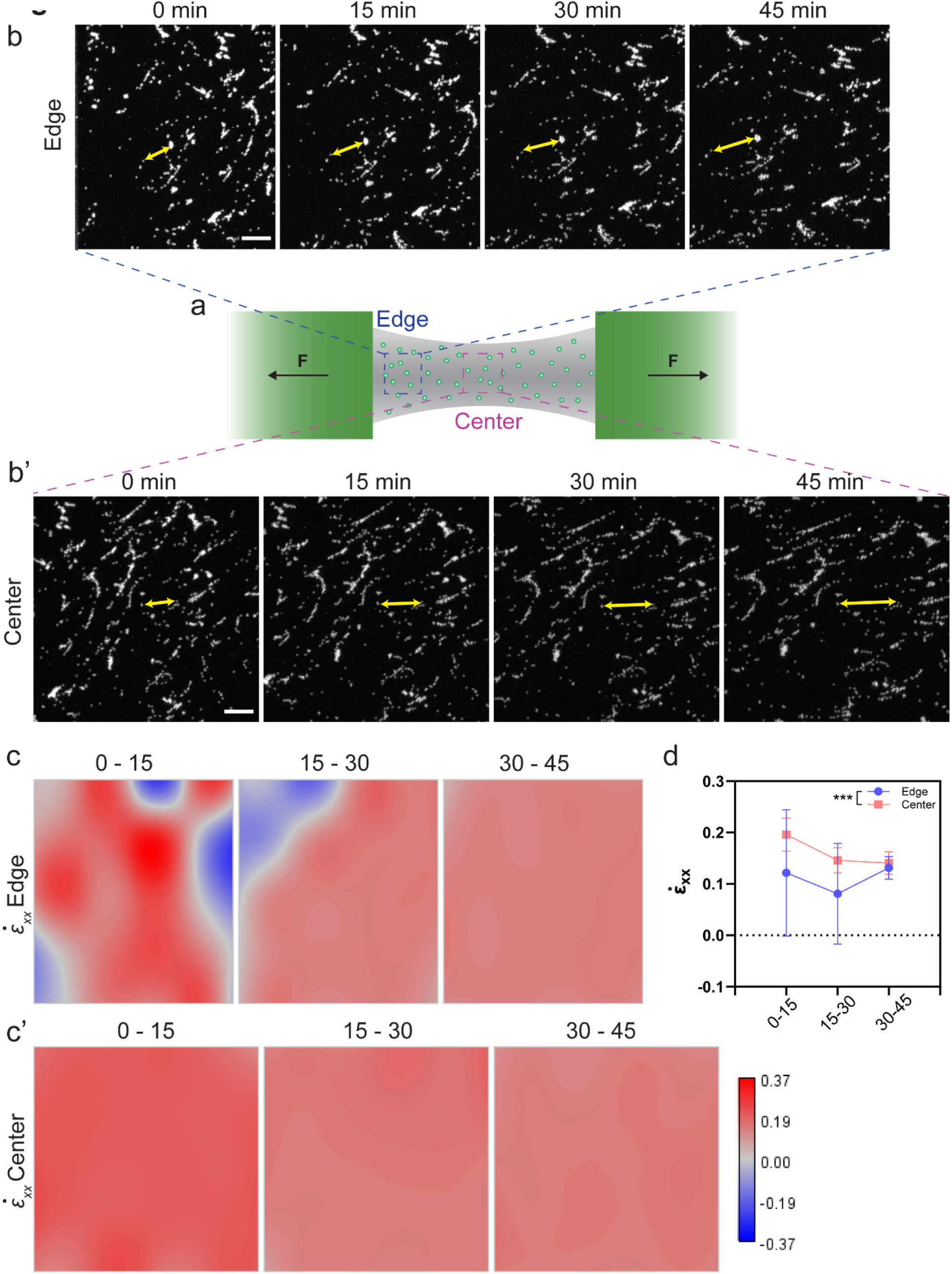
Characterize the strain profile of the PDMS substrate. (**a**) a schematic of the PDMS substrate coated with fluorescent beads. Center (magenta box) and edge (blue box) region are indicated on the schematic. (**b**) Fluorescent beads were traced at the edge and (**b**’) the center locations; yellow arrows showing position changes between two beads. Scale bar = 100 μm (**c**) ε̇_XX_ strain rate map of the edge and (**c**’) the center of the PDMS substrate. A phase lookup table was applied, where blue indicates contraction and red indicates elongation of the material along x-axis. (**d**) Quantification of ε̇_XX_ of edge and center over 0-15, 15-30, 30-45 minutes. Error bars, standard deviation.

To quantify strain uniformity on the PDMS substrate, we used a custom macro for digital image correlation to analyze strain between image pairs across regions of interests (StrainMapper)^3,51^. StrainMapper assumes the image represents a continuous field and carries out a warping transformation^52^ allowing the calculation of principal strain rates ε̇_XX_, ε̇_YY_, and ε̇_XY_. Here we compared strain rate ε̇_XX_ along the x-axis (e.g., the stretch axis) between the edge and the center of the PDMS substrate since the stretcher system is a uniaxial tension system.

Elongation along x-axis results in positive strain whereas shortening results in negative strain, shown in red and blue in the strain rate ε̇_XX_ colormap, respectively (**Figure 2c**). During the initial 30-minute interval, the edge of the PDMS where it is bonded to PES, exhibited heterogeneous strain with large variance (𝜎^2^ = 0.013, 6.2 x 10^-3^ for 0 – 15 min, and 15 – 30 min, respectively) across the field of view. This period was marked by localized contraction proximal to the border of the frame with relatively high elongation at the center of the frame. However, the subsequent span of 30 to 45 minutes after the start of stretching shows a uniform strain distribution (𝜎^2^ = 6.5 x 10^-5^) across the edge region of the stretched PDMS (**Figure 2c**, **and 2d**). In contrast to strains at the edge of the PDMS substrate, the variation of strain was consistently small at the center of the substrate (𝜎^2^ = 5.4 x 10^-5^, 7.6 x 10^-5^, 3.7 x 10^-5^ for 0 to 15 min, 15 to 30 min, and 30 to 45 min, respectively) (**Figure 2c’, and 2d**). The uniaxial strain ε̇_XX_ at the center of the PDMS substrate was homogeneous; thus, tissue samples attached to center will experience consistently homogeneous substrate strain during stretching. For the remainder of this study, we focus on tissues and cells within the center of the PDMS substrate.

### 2.3. Quantify tissue and cellular strain in *Xenopus laevis* organotypic explants

To test the stretcher system on live tissues, preliminary studies of stretching *Xenopus laevis* organotypic explants were conducted and imaged with an inverted brightfield microscope (**Supplementary Video 4**). Then, to visualize cells and test the system with a confocal microscope, we expressed membrane-mNeonGreen mRNAs in the *Xenopus* embryos and cultured them to the early to mid-gastrula stage. Regions of ectoderm expressing mNeonGreen were microsurgically dissected from the embryos. To observe apical cell surfaces during imaging, explants were cultured on the prospective undersurface of fibronectin-coated PDMS bound in a cassette for at least an hour at room temperature. To attach explants to the bottom of the cassette, the fibronectin-coated PDMS-cassette was flipped and placed onto a 3D-printed explant-mounting jig, stabilizing the cassette horizontally for secure mounting of the explant to the PDMS undersurface (**Figure 1f**, **3a, and Methods**). Once the tissue adhered to the PDMS, the cassette was inverted so that the apical face of the organotypic explant faced the objective, to allow tracking of the same group of cells during stretch (**Figure 3b**, **red box: tissue, asterisks: cell**).

**Figure 3.**
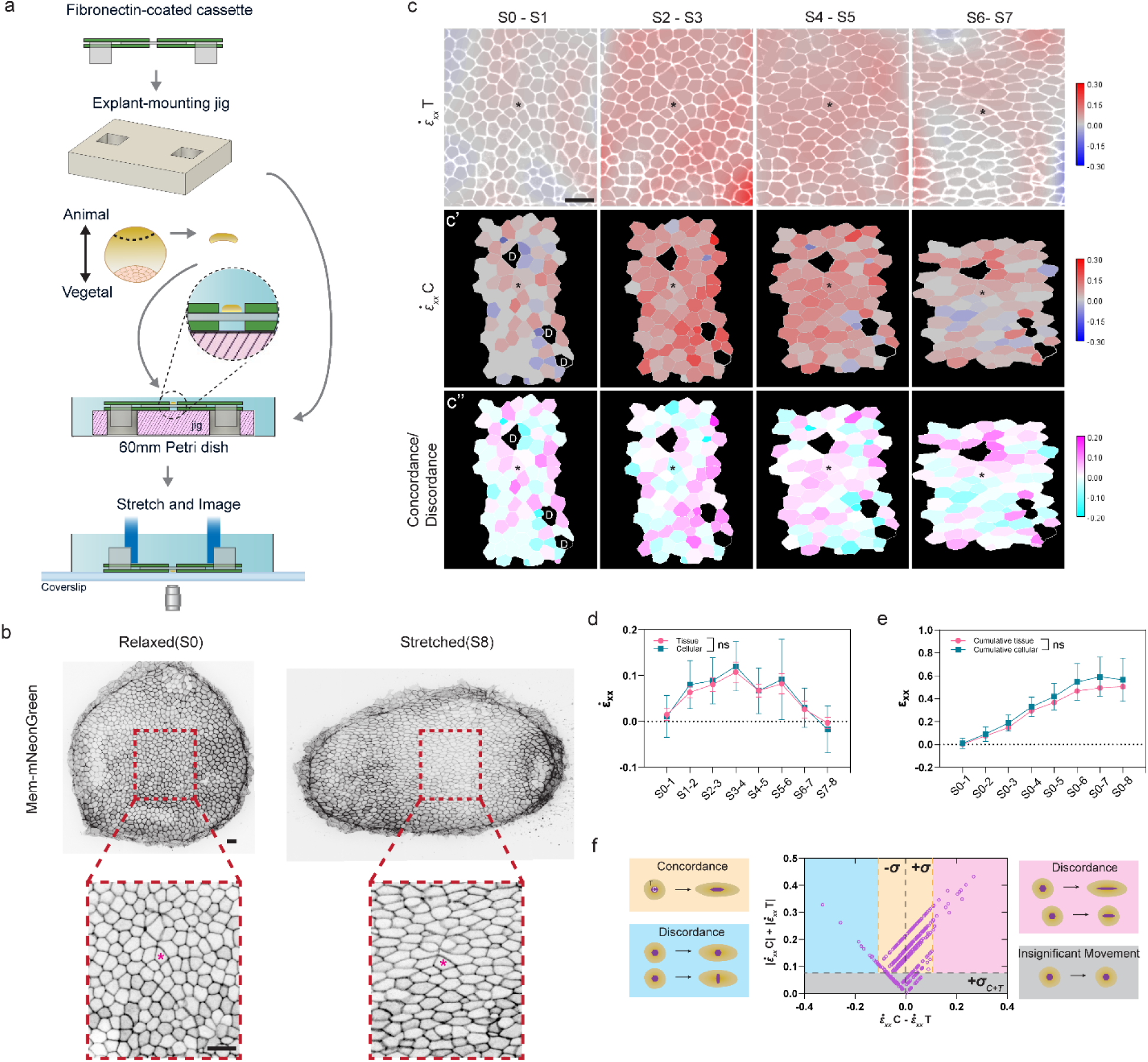
Quantify tissue and cellular strain in *Xenopus laevis* organotypic explants. (**a**) Schematic of mounting *Xenopus laevis* organotypic explant on the fibronectin-coated cassette. An explant-mounting jig was used to provide a flat and stable surface for the flipped cassette. An animal cap organotypic explant was microsurgically removed from a gastrula stage embryo and attached to the PDMS substrate of the cassette and cultured to desired stage. The cassette was flipped back and put into the microscope stage insert for stretch and imaging. (**b**) a Stage 13 animal cap organotypic explant labeled with membrane-mNeonGreen at relaxed state (left) and stretched state (right). Red dashed line outlined the region-of-interests, where the same region was traced and imaged through 8 stretch steps. Scale bar = 35 μm. * indicated the same cell before and after stretching. (**c**) ε̇_XX_𝑇 strain rate map overlayed with cell outlines represented continuous tissue strain rate over two consecutive stretch steps. (**c**’) ε̇_XX_𝐶 strain rate mapped using individual cellular strain, which strain was calculated based on individual cell shape, with no variation of strain within the single cell. N, 81 cells. (**c**’’) A concordance/discordance map was represented by the differences between ε̇_XX_𝐶 and ε̇_XX_𝑇. A magenta-cyan phase lookup table was applied, where both magenta and cyan indicated discordance behaviors between tissue and the individual cell, and white represented concordance. * indicated the same cell across the frames. D represented cells that divided throughout stretching. (**d**) Quantification of both tissue and cellular strain rate ε̇_XX_ between each two consecutive stretch steps. Error bars, standard deviation. (**e**) Quantification of cumulative tissue and cellular strain ε̇_XX_ across the 8 stretch steps. Error bars, standard deviation. (**f**) Tissue-cell concordance/discordance by plotting differences between ε̇_XX_𝐶 and ε̇_XX_𝑇 against absolute values of their sum. Grey region represented insignificant movement of both tissue and the cell. Yellow region represented tissue and the cell strained concordantly. Magenta region represented discordance that the cell strained larger than the tissue. Magenta region represented discordance that the tissue strained larger than the cell. T = tissue; C = cell. 𝜎_C+T_ = 0.081, standard deviation of |ε̇_XX_𝐶| + |ε̇_XX_𝑇|. 𝜎 = 0.057, standard deviation of ε̇_XX_𝐶 - ε̇_XX_𝑇.

To better mimic *in vivo* tissue strain such as those during early development,^1,3,21,33,34,53^ actuators generated up to 150% strain within the cassette. We use the term stretch step (S) to represent a single stretch. To minimize image blurring that occurred during displacement, cell groups were tracked and imaged over 8 additive stretch steps, e.g. S1 to S8 (**Figure 3b**, **and Supplementary Video 5**). In this case, one stretch step displaces cassette ends by 375 μm in total; the initial relaxed state is indicated as S0 and the maximum stretched state, after 8 stretch steps, as S8 (3 mm total grip-to-grip displacement).

To compare tissue and individual cellular mechanical behaviors under tension, we used two independent analysis pipelines to quantify tissue strains and individual cellular strains along the x-axis, roughly along the axis of stretch and represented by subscript *xx*. As before, we used a custom image processing macro (StrainMapper) to calculate tissue strain rate ε̇_XX_𝑇 between two consecutive stretch steps. We found variations in strain within the tissue in each stretch step (σ^2^ ∼ 0.005) (**Figure 3c**, **and 3d**) were greater than variations in the PDMS substrate strain determined above (σ^2^ ∼ 1×10^-5^) (**Figure 2c’, and 2d**). This suggests that the mechanical heterogeneity of the tissues under tension reflects sample variation rather than substrate heterogeneity. From this observation, we suspected that individual cells also experienced heterogenous strain. To measure strain on a cell-by-cell basis, we segmented and registered individual cells across stretch steps with a segmentation software (Seedwater Segmenter^54^). A custom image processing pipeline (FIJI and MATLAB^28,55^) was used to quantify shape and position information from segmented cells. From cell shape changes we calculated the individual cellular strain rate ε̇_XX_𝐶 between two stretch steps of cells that stayed in the imaging frame through all 8 stretch steps. The analysis excluded cells that divided during stretch, since those may not accurately reflect changes in strain that is solely due to the stretching. Similar to our findings from the bulk analysis pipeline, we observed heterogeneous cellular strain (σ^2^ varied from 0.002 to 0.007) across the tissue, with both positively and negatively strained cells (**Figure 3c’**).

We next wanted to investigate how consistently the tissue and individual cells react to tension. Comparing ε̇_XX_𝑇 and ε̇_XX_𝐶 at each stretch step (S0 to S1, S1 to S2, etc.), we found no significant differences (**Figure 3d**). We also compared cumulative tissue and cellular strains ε̇_XX_by calculating strain between the relaxed state (S0) and each stretch step (S1, S2, etc.). Again, we found no significant differences between cumulative tissue and cellular strain at each stretch step (**Figure 3e**). To investigate cell-to-cell variation between tissue and cell strain, we mapped differences between the two analytic pipelines across the tissue reporting whether tissue and cell strains were in *concordance* or *discordance* (**Figure 3c’’**). The cell and tissue strains are in concordance if the difference between tissue and cellular strain is within one standard deviation. If the difference is greater than one standard deviation, the cellular and tissue strain are in discordance. We observed most tissue-cell strains were concordant (**Figure 3f**, **yellow region**), suggesting consistent tissue and cell behaviors under tension. We also observed cases where (1) tissues and cells only displayed small fluctuations in strain (**Figure 3f**, **grey region**), (2) individual cells strained more than the tissue, resulting in discordant strain (**Figure 3f**, **magenta region**), and (3) individual cell strains that were less than the tissue or even contracted while the tissue was stretched (**Figure 3f**, **cyan region**). Thus, we find that strain is heterogeneously distributed through *Xenopus* organotypic animal cap explant when under tension, with individual cells that showed either concordant, or discordant strains within the tissue. Combining our novel stretcher with image analysis pipelines reveals more complex behaviors of cells to strain than previously seen.

### 2.4. High-resolution live imaging of various model systems on the stretcher

Building on the previous demonstration, we next sought to use the stretcher system with high resolution confocal fluorescent live-imaging, testing the compatibility of the cassette with a high numerical-aperture oil-immersion objective lens. We stretched *Xenopus* organotypic explants that expressing an intermediate filament reporter, keratin8-mCherry, and a membrane marker, membrane-mNeonGreen. We were able to observe and track keratin filament changes at a single filament level using a high numerical-aperture oil-immersion objective lens (**Figure 4a**). Prior to stretch, we observed many curved keratin filaments suggesting a relaxed state.

**Figure 4.**
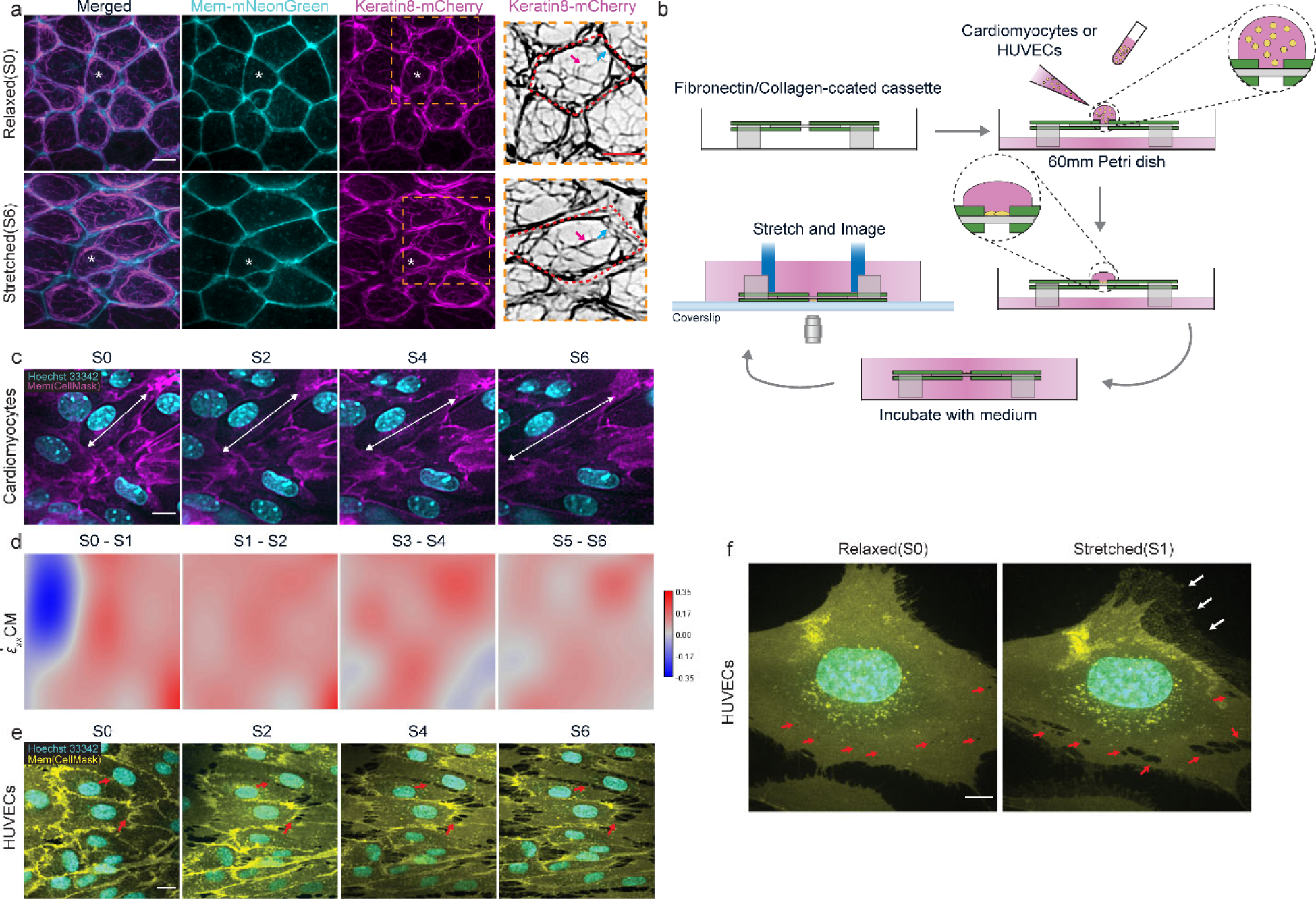
High-resolution live imaging of various model systems on the stretcher. (**a**) A Stage 11 animal cap organotypic explant labeled with mem-mNeonGreen and keratin8-mCherry at relaxed state (top) and stretched state (bottom). * indicates the same cell across frames. The last column shows keratin filaments in a single cell. Red dashed line outlined the single cell. Magenta arrow highlighted the filament that was tortuous at relaxed state but straightened after stretching. Cyan arrow highlighted the separation between the filaments and the cell junction. Scale bar = 10 μm. (**b**) Schematic of seeding cardiomyocytes or HUVECs on fibronectin- or collagen-coated cassette. A droplet of suspended cardiomyocytes or HUVECs was pipetted onto the PDMS substrate of the cassette with 500 μL of respective medium at the bottom of the petri dish. After cells attach to the PDMS substrate, 12 mL of medium was added to fully submerge the cassette. The cassette was flipped back and put into microscope stage insert before imaging. (**c**) Cardiomyocytes labeled with membrane and nucleus markers were stretched and imaged for 6 stretch steps. White arrows indicated straining of the cell membrane. Scale bar = 10 μm. (**d**) ε̇_XX_𝐶𝑀 strain rate map of cardiomyocytes. (**e**) HUVECs labeled with membrane and nucleus markers were stretched and imaged for 6 stretch steps. Red arrows highlighted tearing of cell-cell contacts. Scale bar = 20 μm. (**f**) a single HUVEC at relaxed (left) and stretched (right) state. Red arrows highlighted the fenestrated holes in the cell membrane. White arrows highlighted the detachment of cell membrane to the PDMS substrate. Scale bar = 10 μm.

However, filaments straightened as stretch increased, suggesting keratin filaments bear more load after stretching (**Figure 4a**, **red arrows**). We also observed some cases where keratin filaments initially associated with the cell-cell junction would, after stretching, detach from junctions (**Figure 4a**, **blue arrows**). Both cases of keratin network remodeling support their role in carrying loads under tension.^56,57^

We also tested two other live cell model systems in our stretcher system, mouse neonatal cardiomyocytes and human umbilical vein endothelial cells (HUVECs). To seed the cells on the cassette, as before, we prepared ECM-coated cassettes before the experiment by adsorbing either fibronectin (HUVECs) or collagen I (cardiomyocytes) onto the PDMS substrates. After coating, we inverted the cassette and loaded cells onto the PDMS in 30 uL of medium containing either cardiomyocytes or HUVECs. We incubated cells on the cassette until they attached to the PDMS surface. Vital dyes for nuclei and plasma membrane were added to the medium 30 minutes before imaging. As with tissue explants, we inverted the cassette, and placed it in the stretcher (**Figure 4b**, **and Methods**). We applied strain and collected images over 6 stretch steps.

Cardiomyocytes appeared to maintain cell-cell contacts while nuclei shape remained unchanged as they were stretched (**Figure 4c**). A consistent field of cells throughout the stretch steps allowed us to map tissue strain between each consecutive stretch steps ε̇_XX_𝐶𝑀 using StrainMapper (**Figure 4d**). Cardiomyocytes under high strain remain connected to each other and continue to beat (**Supplementary Video 6**).

Since HUVECs are considerably larger than Xenopus embryonic cells or cardiomyocytes, we initially used a 25x objective to image confluent layers to include more cells in the imaging frame. In contrast to our *Xenopus* embryonic epithelium and mouse cardiomyocytes, applied strain disrupted HUVECs cell-cell adhesions, and caused cell membranes to detach from the PDMS substrate (**Figure 4e**). Since stretching caused HUVEC membranes to retract and expose gaps between cells, we were not able to perform a tissue strain analysis. Next, we used a higher numerical aperture objective and observed single isolated cells under high substrate strain. We lowered the seeding density, allowing us to strain single HUVECs (**Figure 4f**). While not obvious at rest, single cells under high strain exhibited multiple holes, similar to fenestrations that had been described previously in other endothelial tissues.^58^ At increased strains, cell membranes appeared to rupture or tear at the where fenestrated holes had appeared at lower strains (**Figure 4f**, **red arrows**). Furthermore, cell-substrate adhesion was also disrupted after stretching, leaving only streaks of membrane attached to the substrate (**Figure 4f**, **white arrows**). Comparing these two cell types, our preliminary observations indicate cardiomyocytes can remain attached to the substrate and maintain cell-cell contacts under strain. By contrast, highly strained confluent layers of HUVECs lost cell-cell adhesions and single cells ruptured at sites where fenestrations were observed, and both single, and confluent HUVECs lost cell-substrate adhesions.

## 3. Discussion

In this work, we developed a stretcher system, the TissueTractor, that enables an effective mechanical stimulation of live samples while achieving high resolution live imaging. The stretcher makes use of affordable 2D cutters and 3D printers to fabricate parts that can be integrated with low-cost linear actuators. Key to the stretcher system is an exchangeable 2D-cut cassette that keeps the sample within the working distance of a high numerical-aperture objective lens. The cassette and two linear actuators are assembled into a custom designed microscope stage insert for simultaneous mechanical manipulation and visualization. With the stretcher system, we were able to acquire high-resolution live images of organotypic explants during stretching and analyze tissue and cellular scale engineering strain. In addition, we were able to visualize intracellular intermediate filaments under strain at a single filament level. We also stretched cardiomyocytes isolated from neonatal mice and endothelial cells from human umbilical vein. These model tissues demonstrate the utility of the exchangeable cassettes and our stretcher system for visualizing and measuring the effects of strain on diverse cell and tissue types under strain with high-resolution confocal imaging. Although we only tested three animal models in this work, the stretcher system is compatible with most cultured cells, and organotypic explants that adhere to extracellular matrix. Furthermore, the low-cost, lightweight, and modular features of the stretcher system provide extensive customization for specific applications.

Various adaptations can be easily integrated to broaden the applications of the stretcher system. The cassette can be adapted to use other biocompatible adhesives to attach tissue samples to PDMS substrates such as cyanoacrylate, Cell-Tak (Corning), or poly-L-lysine.^5,59^ Furthermore, in the future we envision modifying the cassette design by changing grip positions or spring connections to provide different strain profiles (i.e. shear strain, biaxial strain) (**Figure S5**), to mimic other dynamically changing microenvironments. Furthermore, the size of the cassette substrate can be increased to include larger numbers of tissue/cell samples for fixation. While the current stretcher design sought to track and visualize live cellular and intracellular dynamics during stretching, other systems with multiple coupled stretchers may achieve high-throughput analysis of mechano-responses.

Our stretcher system will enable testing of putative mechanosensors and mechanotransducers in living cells and provide insights into the signaling pathways and gene regulatory networks that respond to mechanical stimulation. In particular, the ability to generate and sustain high strain rates make this a unique tool to investigate the plastic behaviors of growing multicellular tissues and how mechanical cues play a role in development and disease.

## 4. Methods

### Preparation and fabrication of the stretcher system

The top and bottom microscope stage insert, and the H-bridge were printed with Polylactic Acid (PLA; Prusa Research) filaments in Fused Deposition Modeling 3d-printer (Prusa i3 MK3S; Prusa Research). Polyester (PES) sheets were purchased from Precision Brand.

Poly(dimethysiloxane) (PDMS) sheets (0.005’’, 40D, Gloss finish) were purchased from Specialty Manufacturing, inc. Top and bottom PES sheets of the cassette, and the dumbbell-shape PDMS sheets were cut using Roland CAMM-1 GS-24 Vinyl Cutter (Roland DGA) with a 25 degree/.125 offset blade (USA-C125; Roland DGA) to provide clean, smooth cutting edges. Then, PES and PDMS sheets were washed with 100% acetone, followed by extensive rinse of 100% ethanol and double deionized water for 24 hours. Rinsed products were dried between wax paper to prevent dust accumulation. Stretcher blocks were 3D-printed using stereolithography (Form 2; Formlabs) with photocurable clear resin (Formlabs), followed by a 20-minute 100% isopropanol wash (Form Wash; Formlabs) to remove excess resin, and then cured in UV-light (405 nm) chamber (Form Cure; Formlabs) for 2 hours at 60°C.

The bottom PES sheets were placed in a 3D-printed jig to facilitate assembly (**Figure S3**). The dumbbell-shape PDMS sheet was placed onto the bottom PES sheet with the dumbbell parts aligned with the protruding parts of the sheet. A thin layer of UV-curable optical adhesive (Norland Optical Adhesive 63; Edmund Optics) was applied to the PDMS and the PES sheet (excluding the spring-like structures). Then, top sheets were placed onto the bottom sheet using the jig. Two stretcher blocks were bonded into the two cut-out holes on the top sheets (**Figure 1c’, and Figure S3**). Once assembled, the cassette was placed into UV-light (350nm) chamber to cure for 2 hours. Cassettes were stored at room temperature before use and discarded after experiments.

### Fluorescent bead coating

30 µL of green fluorescent polymer microspheres (5.0 µm diameter; 1% solids; Duke Scientific Corp.) was diluted in 420 µL of double deionized water. A cassette was flipped (stretcher blocks at the bottom) and 100 µL of the diluted fluorescent beads solution was added onto the PDMS part of the cassette. The cassette was air-dried, covered with aluminum foil, until the liquid evaporated.

### Microinjection of *Xenopus laevis* embryos and organotypic explant mounting

All *Xenopus laevis* work was approved by the University of Pittsburgh Division of Laboratory Animal Resources. *Xenopus laevis* embryos were obtained by the standard procedure.^60^ 50 pg of membrane-tagged mNeonGreen mRNA was microinjected into 4-cell embryos. To visualize keratin 8 filaments, 100 pg of keratin8-mCherry mRNA was microinjected into 4-cell embryos. Injected and uninjected embryos were cultured in 1/3x modified Barth Solution (MBS)^61^ to desired stages.

Cassettes were flipped (stretcher blocks at the bottom) and put into an oxygen plasma cleaner (Harrick Plasma) for 2 minutes to activate surface of the PDMS substrates. 0.025 µg/µl of fibronectin (Chem Cruz) was added onto the cassette immediately after plasma cleaning and incubated at room temperature for 1 hour. The cassette was then transferred into a 60 mm petri dish with a mounting jig in Danilchik’s For Amy^61^ medium with antibiotic and antimycotic (Sigma) (**Figure 3a**). An organotypic animal cap explant was microsurgically removed^62^ at early gastrula stage (Stage 10)^63^ and immediately transferred and positioned at the center of the PDMS substrate of the cassette (**Figure 3a**). A 1.5 mm x 12 mm glass coverslip bridge with high vacuum grease (DuPont) on both ends was gently pressed down to immobilize the explant and incubated at 14°C or room temperature to the desired stage. The glass bridge was removed before the cassette was transferred to the stretcher.

### Human umbilical vein endothelial cell (HUVECs) culture

Pooled human umbilical vein endothelial cells (HUVECs; Promocell) were cultured in a sterile humidified incubator in complete endothelial cell growth medium (EC Growth Medium 2/EGM2, containing 2% FBS; Promocell) and 1× antibiotic-antimycotic (Thermo Fisher Scientific) at 37 °C under 5% CO2. Upon confluency, cells were rinsed with HEPES BSS (Detach Kit, Promocell) and treated with 0.04% trypsin/0.03% EDTA for 5-7 minutes at room temperature until cells detached. After adding trypsin neutralization solution, cells were centrifuged at 220 × g for 3 minutes and gently resuspended in fresh EGM2. Cells were maintained in EGM2 for a maximum of six passages.

### Mouse neonatal cardiomyocyte isolation and culture

All mouse work was approved by the University of Pittsburgh Division of Laboratory Animal Resources. Outbred Swiss Webster mice were used to generate cardiomyocytes for stretcher experiments. Neonatal mouse cardiomyocytes were isolated as described.^64^ Briefly, mouse pups were sacrificed at P2 and the hearts were removed, cleaned, minced, and digested overnight at 4°C in 20 mM BDM (2,3-butanedione monoxime) and 0.0125% trypsin in Hank’s balanced salt solution. The next day, heart tissue was digested further in 15 mg/ml Collagenase/Dispase (Roche) in Leibovitz medium with 20 mM BDM to create a single-cell suspension. Cells were preplated for 1.5–2 h in plating medium (65% high glucose DMEM, 19% M-199, 10% horse serum, 5% fetal bovine serum, and 1% penicillin-streptomycin) to remove fibroblasts and endothelial cells. After preplating, cardiomyocytes were counted manually on a hemocytometer and cell density was adjusted to 3,000,000 cells/mL to seed onto the stretcher.

### HUVECs and mouse cardiomyocytes seeding and live-labeling

Cassettes were flipped (stretcher blocks at the bottom) and two 20 mm x 10 mm PDMS sheets were placed onto the PES sheet part of the cassette, exposing the PDMS substrate. This ensured that only the PDMS substrate was surface activated in the oxygen plasma cleaner (Harrick Plasma) and the PES sheet remained hydrophobic. The cassettes were surface activated in the oxygen plasma cleaner for 2 minutes. 0.025 µg/µl of fibronectin (Chem Cruz) or 0.25 µg/µl of rat tail Type I collagen (Millipore) were added onto the cassette immediately after plasma cleaning, followed by 1-hour incubation at room temperature. The fibronectin-coated cassettes were then stored at 4°C until use. Collagen Type I was aspirated out from the surface of the cassette, followed by 1-hour UV curing. Then the collagen-coated cassettes were washed using phosphate-buffered saline (PBS; Sigma) and stored dry, covered with aluminum foil, at room temperature until use.

The 20 mm x 10 mm PDMS sheets were removed from the cassette before seeding. 30 µL of HUVECs (250,000 cells/mL) or mouse cardiomyocytes (3,000,000 cells/mL) were placed onto the PDMS part of the fibronectin-coated or collagen I-coated cassette, respectively, to form a droplet on top of the PDMS substrate (**Figure 4b**).

Cassettes with HUVECs droplets were cultured at 37°C for 2 hours to allow cell attachment, and then 8 mL of complete growth medium (Endothelial Cell Basal Medium-2 C-22211, with Endothelial Cell Growth Medium 2 Supplement Pack C-39211, Promocell) was added into the petri dish to fully submerge the cassette. Submerged cassettes were incubated for 16 hours at 37°C before imaging.

Cassettes with cardiomyocytes were cultured at 37°C for 4 hours to allow cell attachment. Due to the long incubation time, 500 µL of cardiomyocyte plating medium was added to the bottom of the petri-dish to limit evaporation during incubation. After 4 hours, 12 mL of cardiomyocyte plating medium was added to the dish to fully submerge the cassette. 16 hours post-plating, the plating medium was exchanged for cardiomyocyte maintenance medium (78% high glucose DMEM, 17% M-199, 4% horse serum, 1% penicillin/streptomycin, 1 μm Ara-C, and 1 μm isoproterenol) and incubated at 37°C for 72 hours before imaging.

To label cells for live cell imaging, cassettes were flipped back (stretcher blocks at the top) and Hoechst 33342 (nuclear stain; 2 µg/mL; Thermo Fisher Scientific) and CellMask Green (plasma membrane stain, 1x working solution; Thermo Fisher Scientific) were added to the medium. Cells were incubated at 37°C in 5% CO_2_ for 30 minutes in stain prior to imaging.

### Stretcher system setup and Imaging

The cassette was transferred and positioned at the center of the sample chamber of the stage bottom with 4 mL of respective medium. Then, the stage top was placed onto the stage bottom; two picomotor piezo linear actuators (8301NF; Newport) with fully extended arms were securely mounted at the motor mount. The H-bridge was then inserted until the two ends fully contacted the H-bridge rest of the stage top (**Figure 1e**). The assembled microscope stage insert was placed onto an inverted microscope (**Figure S1**).

The picomotor piezo linear actuators can be controlled by a LabView program with customizable velocity settings or manually operated by a joystick. Images at relaxed state and at the end of each stretch step were acquired using an inverted compound microscope (Leica) with a 63x/1.40NA oil immersion or a 25x/0.95NA water immersion objective lens, equipped with a spinning disk scanhead (Yokogawa) and a CMOS camera (Hamamatsu). Sequential images were acquired using a microscope automation software (μManager 2.0)^50^.

### Segmentation and strain analysis

Seedwater Segmenter^54^ was used to segment *Xenopus* epithelial cells. A custom FIJI macro was used to acquire cell ROIs and shape information from the segmented cells. PDMS deformation and tissue level strains were calculated using beta-spline based image registration (ImageJ plugin bUnWarpJ)^52^ and a custom ImageJ macro (StrainMapper); custom MATLAB m-code calculated cell level engineering strain rates.^55^ Custom macros and codes are available upon request.

## Supporting Information

Supporting Information is available from the author.

## Acknowledgements

We would like to thank Geneva Masak, and other members of the Davidson group for their comments and discussions. We thank Drs. Sagar Joshi, Rafey Feroze, and Steven Trier for their early work on prototype stretchers in the lab. This work was supported by National Institutes of Health (R37 HD044750). E.H. was additionally supported by a Summer Research Internship from the Swanson School of Engineering. S.B. was supported by an American Heart Association Predoctoral Fellowship. A.K. was supported by the National Institutes of Health (R01 HL127711). T.W. was supported by the National Science Foundation NSF Systems Medicine REU (EEC #1156899). Y.C. was also supported by National Institutes of Health (R01 HL136566) and the Berenfield Graduate Fellowship in Bioengineering. S.A. and D.S.V. were additionally supported by the NIH Biomechanics in Regenerative Medicine Training Grant (BiRM; T32 EB003392). We thank Dr. Viki Allan for sharing the *Xenopus laevis* eGFP-Keratin8.1S plasmid.

## Author contributions

Conceptualization: J.Y., E.H., D.V., T.W., and L.D.; Methodology: J.Y., and L.D.; Software: J.Y. D.V., S.A., and L.D.; Formal Analysis: J.Y., and L.D.; Investigation: J.Y., Y.C., and S.B.; Resources: J.Y., E.H., Y.W., D.V., C.S., A.K., and L.D.; Writing – Original Draft: J.Y., and L.D.; Writing – Review & Editing: J.Y., E.H., Y.W., D.V., T.W., S.A., C.S., Y.C., S.B., A.K., and L.D.; Visualization: J.Y.; Supervision: L.D.; Funding Acquisition: A.K., and L.D.

## Data Availability

Source data are provided with this paper. All other data that support the findings of this study are available from the authors upon reasonable request.

## Ethics approval

All *Xenopus laevis* and mouse work were approved by the Institutional Animal Care and Use Committee (Protocol #: 24014521) and the University of Pittsburgh Division of Laboratory Animal Resources.

## References

1 Davidson, L., von Dassow, M. & Zhou, J. Multi-scale mechanics from molecules to morphogenesis. Int J Biochem Cell Biol 41, 2147–2162, doi:10.1016/j.biocel.2009.04.015 (2009).

2 Engler, A. J., Sen, S., Sweeney, H. L. & Discher, D. E. Matrix elasticity directs stem cell lineage specification. Cell 126, 677–689, doi:10.1016/j.cell.2006.06.044 (2006).

3 Feroze, R., Shawky, J. H., von Dassow, M. & Davidson, L. A. Mechanics of blastopore closure during amphibian gastrulation. Dev Biol 398, 57–67, doi:10.1016/j.ydbio.2014.11.011 (2015).

4 Gjorevski, N. & Nelson, C. M. The mechanics of development: Models and methods for tissue morphogenesis. Birth Defects Res C Embryo Today 90, 193–202, doi:10.1002/bdrc.20185 (2010).

5 Schluck, T., Nienhaus, U., Aegerter-Wilmsen, T. & Aegerter, C. M. Mechanical control of organ size in the development of the Drosophila wing disc. PLoS One 8, e76171, doi:10.1371/journal.pone.0076171 (2013).

6 Heer, N. C. & Martin, A. C. Tension, contraction and tissue morphogenesis. Development 144, 4249–4260, doi:10.1242/dev.151282 (2017).

7 Ingber, D. E., Prusty, D., Sun, Z., Betensky, H. & Wang, N. Cell shape, cytoskeletal mechanics, and cell cycle control in angiogenesis. J Biomech 28, 1471–1484, doi:10.1016/0021-9290(95)00095-x (1995).

8 Jackson, T. R., Kim, H. Y., Balakrishnan, U. L., Stuckenholz, C. & Davidson, L. A. Spatiotemporally Controlled Mechanical Cues Drive Progenitor Mesenchymal-to-Epithelial Transition Enabling Proper Heart Formation and Function. Curr Biol 27, 1326–1335, doi:10.1016/j.cub.2017.03.065 (2017).

9 Kim, H. Y., Jackson, T. R., Stuckenholz, C. & Davidson, L. A. Tissue mechanics drives regeneration of a mucociliated epidermis on the surface of Xenopus embryonic aggregates. Nat Commun 11, 665, doi:10.1038/s41467-020-14385-y (2020).

10 Zorn, A. M. & Wells, J. M. Vertebrate endoderm development and organ formation. Annu Rev Cell Dev Biol 25, 221–251, doi:10.1146/annurev.cellbio.042308.113344 (2009).

11 Northcott, J. M., Dean, I. S., Mouw, J. K. & Weaver, V. M. Feeling Stress: The Mechanics of Cancer Progression and Aggression. Front Cell Dev Biol 6, 17, doi:10.3389/fcell.2018.00017 (2018).

12 Thompson, A. J. et al. Rapid changes in tissue mechanics regulate cell behaviour in the developing embryonic brain. Elife 8, doi:10.7554/eLife.39356 (2019).

13 Chu, C., Masak, G., Yang, J. & Davidson, L. From biomechanics to mechanobiology: Xenopus provides direct access to the physical principles that shape the embryo. Current Opinion in Genetics & Development 63, 71–77 (2020).

14 Argentati, C. et al. Insight into Mechanobiology: How Stem Cells Feel Mechanical Forces and Orchestrate Biological Functions. Int J Mol Sci 20, doi:10.3390/ijms20215337 (2019).

15 Petridou, N. I., Spiro, Z. & Heisenberg, C. P. Multiscale force sensing in development. Nat Cell Biol 19, 581–588, doi:10.1038/ncb3524 (2017).

16 O’Connor, J. W. & Gomez, E. W. Biomechanics of TGFbeta-induced epithelial-mesenchymal transition: implications for fibrosis and cancer. Clin Transl Med 3, 23, doi:10.1186/2001-1326-3-23 (2014).

17 Koser, D. E. et al. Mechanosensing is critical for axon growth in the developing brain. Nat Neurosci 19, 1592–1598, doi:10.1038/nn.4394 (2016).

18 Takao, S., Taya, M. & Chiew, C. Mechanical stress-induced cell death in breast cancer cells. Biol Open 8, doi:10.1242/bio.043133 (2019).

19 Barriga, E. H., Franze, K., Charras, G. & Mayor, R. Tissue stiffening coordinates morphogenesis by triggering collective cell migration in vivo. Nature 554, 523, doi:10.1038/nature25742 (2018).

20 Sadler, T. W. & Feldkamp, M. L. The embryology of body wall closure: relevance to gastroschisis and other ventral body wall defects. Am J Med Genet C Semin Med Genet 148C, 180–185, doi:10.1002/ajmg.c.30176 (2008).

21 Chien, Y. H., Keller, R., Kintner, C. & Shook, D. R. Mechanical strain determines the axis of planar polarity in ciliated epithelia. Curr Biol 25, 2774–2784, doi:10.1016/j.cub.2015.09.015 (2015).

22 Mongera, A. et al. A fluid-to-solid jamming transition underlies vertebrate body axis elongation. Nature 561, 401–405, doi:10.1038/s41586-018-0479-2 (2018).

23 Bi, D., Yang, X., Marchetti, M. C. & Manning, M. L. Motility-driven glass and jamming transitions in biological tissues. Phys Rev X 6, doi:10.1103/PhysRevX.6.021011 (2016).

24 Atia, L., Fredberg, J. J., Gov, N. S. & Pegoraro, A. F. Are cell jamming and unjamming essential in tissue development? Cells Dev 168, 203727, doi:10.1016/j.cdev.2021.203727 (2021).

25 Trepat, X. et al. Universal physical responses to stretch in the living cell. Nature 447, 592–595, doi:10.1038/nature05824 (2007).

26 Peifer, M. et al. Polychaetoid/ZO-1 strengthens cell junctions under tension while localizing differently than core adherens junction proteins. Mol Biol Cell 34, ar81, doi:10.1091/mbc.E23-03-0077 (2023).

27 Banavar, S. P. et al. Mechanical control of tissue shape and morphogenetic flows during vertebrate body axis elongation. Sci Rep 11, 8591, doi:10.1038/s41598-021-87672-3 (2021).

28 Sasai, Y. Cytosystems dynamics in self-organization of tissue architecture. Nature 493, 318–326, doi:10.1038/nature11859 (2013).

29 Eiraku, M. et al. Self-organizing optic-cup morphogenesis in three-dimensional culture. Nature 472, 51–56, doi:10.1038/nature09941 (2011).

30 Palmquist, K. H. et al. Reciprocal cell-ECM dynamics generate supracellular fluidity underlying spontaneous follicle patterning. Cell 185, 1960–1973. e1911 (2022).

31 Shyer, A. E., Huycke, T. R., Lee, C., Mahadevan, L. & Tabin, C. J. Bending gradients: how the intestinal stem cell gets its home. Cell 161, 569–580, doi:10.1016/j.cell.2015.03.041 (2015).

32 Shyer, A. E. et al. Emergent cellular self-organization and mechanosensation initiate follicle pattern in the avian skin. Science 357, 811–815 (2017).

33 Glickman, N. S., Kimmel, C. B., Jones, M. A. & Adams, R. J. Shaping the zebrafish notochord. Development 130, 873–887, doi:10.1242/dev.00314 (2003).

34 Zhou, J., Pal, S., Maiti, S. & Davidson, L. A. Force production and mechanical accommodation during convergent extension. Development 142, 692–701, doi:10.1242/dev.116533 (2015).

35 Vijayraghavan, D. S. & Davidson, L. A. Mechanics of neurulation: From classical to current perspectives on the physical mechanics that shape, fold, and form the neural tube. Birth Defects Res 109, 153–168, doi:10.1002/bdra.23557 (2017).

36 Keller, R. & Sutherland, A. in Current Topics in Developmental Biology Vol. 136 271-317 (Elsevier, 2020).

37 Constantinou, I. & Bastounis, E. E. Cell-stretching devices: advances and challenges in biomedical research and live-cell imaging. Trends Biotechnol 41, 939–950, doi:10.1016/j.tibtech.2022.12.009 (2023).

38 Harris, A. R. et al. Generating suspended cell monolayers for mechanobiological studies. Nat Protoc 8, 2516–2530, doi:10.1038/nprot.2013.151 (2013).

39 Wiebe, C. & Brodland, G. W. Tensile properties of embryonic epithelia measured using a novel instrument. J Biomech 38, 2087–2094 (2005).

40 Nestor-Bergmann, A. et al. Decoupling the Roles of Cell Shape and Mechanical Stress in Orienting and Cueing Epithelial Mitosis. Cell Rep 26, 2088–2100 e2084, doi:10.1016/j.celrep.2019.01.102 (2019).

41 Estermann, S. J., Pahr, D. H. & Reisinger, A. Hyperelastic and viscoelastic characterization of hepatic tissue under uniaxial tension in time and frequency domain. J Mech Behav Biomed Mater 112, 104038, doi:10.1016/j.jmbbm.2020.104038 (2020).

42 Duda, M. et al. Polarization of Myosin II Refines Tissue Material Properties to Buffer Mechanical Stress. Dev Cell 48, 245–260 e247, doi:10.1016/j.devcel.2018.12.020 (2019).

43 Sumi, A. et al. Adherens Junction Length during Tissue Contraction Is Controlled by the Mechanosensitive Activity of Actomyosin and Junctional Recycling. Dev Cell 47, 453–463 e453, doi:10.1016/j.devcel.2018.10.025 (2018).

44 Farge, E. Mechanical induction of Twist in the Drosophila foregut/stomodeal primordium. Curr Biol 13, 1365–1377 (2003).

45 Sotoudeh, M., Jalali, S., Usami, S., Shyy, J. Y. & Chien, S. A strain device imposing dynamic and uniform equi-biaxial strain to cultured cells. Ann Biomed Eng 26, 181–189, doi:10.1114/1.88 (1998).

46 Beloussov, L. V., Lakirev, A. V. & Naumidi, I. I. The role of external tensions in differentiation of Xenopus laevis embryonic tissues. Cell Differentiation and Development 25(3), 165–176 (1988).

47 CellScale. <cellscale.com>

48 FLEXCELL. <flexcellint.com>

49 Liu, R. et al. Analysis, simulation and fabrication of MEMS springs for a micro-tensile system. J Micromech Microeng 19, doi:Artn 01502710.1088/0960-1317/19/1/015027 (2009).

50 Edelstein, A. D. et al. Advanced methods of microscope control using μManager software. Journal of biological methods 1 (2014).

51 Stepien, T. L., Lynch, H. E., Yancey, S. X., Dempsey, L. & Davidson, L. A. Using a continuum model to decipher the mechanics of embryonic tissue spreading from time-lapse image sequences: An approximate Bayesian computation approach. PLoS One 14, 460774, doi:10.1371/journal.pone.0218021 (2019).

52 Arganda-Carreras, I. et al. Consistent and elastic registration of histological sections using vector-spline regularization. CVAMIA:Computer Vision Approaches to Medical Image Analysis: 4241, 85–95 (2006).

53 Bhattacharya, D., Zhong, J., Tavakoli, S., Kabla, A. & Matsudaira, P. Strain maps characterize the symmetry of convergence and extension patterns during zebrafish gastrulation. Sci Rep 11, 19357, doi:10.1038/s41598-021-98233-z (2021).

54 Mashburn, D. N., Lynch, H. E., Ma, X. & Hutson, M. S. Enabling user-guided segmentation and tracking of surface-labeled cells in time-lapse image sets of living tissues. Cytometry A 81, 409–418, doi:10.1002/cyto.a.22034 (2012).

55 Vijayraghavan, D. Elucidating the Role of Mechanics in Neural Plate Convergent Extension PhD thesis, University of Pittsburgh, (2018).

56 Sonavane, P. R. et al. Mechanical and signaling roles for keratin intermediate filaments in the assembly and morphogenesis of Xenopus mesendoderm tissue at gastrulation. Development 144, 4363–4376, doi:10.1242/dev.155200 (2017).

57 Broussard, J. A. et al. Scaling up single-cell mechanics to multicellular tissues - the role of the intermediate filament-desmosome network. J Cell Sci 133, jcs228031, doi:10.1242/jcs.228031 (2020).

58 Palade, G. E., Simionescu, M. & Simionescu, N. Structural aspects of the permeability of the microvascular endothelium. Acta Physiol Scand Suppl 463, 11–32 (1979).

59 Vogt, E. J. et al. Author Correction: Anchoring cortical granules in the cortex ensures trafficking to the plasma membrane for post-fertilization exocytosis. Nat Commun 10, 2926, doi:10.1038/s41467-019-10935-1 (2019).

60 Kay, B. K. Xenopus laevis: Practical uses in cell and molecular biology. Injections of oocytes and embryos. Methods Cell Biol 36, 663–669 (1991).

61 Sive, H. L., Grainger, R. M. & Harland, R. M. Early development of Xenopus laevis : a laboratory manual. (Cold Spring Harbor Laboratory Press, 2000).

62 Joshi, S. D. & Davidson, L. A. Live-cell imaging and quantitative analysis of embryonic epithelial cells in Xenopus laevis. J Vis Exp, doi:10.3791/1949 (2010).

63 Nieuwkoop, P. D. & Faber, J. Normal tables of Xenopus laevis (Daudin). (Amsterdam: Elsevier North-Holland Biomedical Press, 1967).

64 Ehler, E., Moore-Morris, T. & Lange, S. Isolation and culture of neonatal mouse cardiomyocytes. J Vis Exp, doi:10.3791/50154 (2013).

