## Supplementary Materials for "The TissueTractor, a device for applying large strains to tissues and cells for simultaneous high-resolution live cell microscopy"

**Supplementary Figure 1**

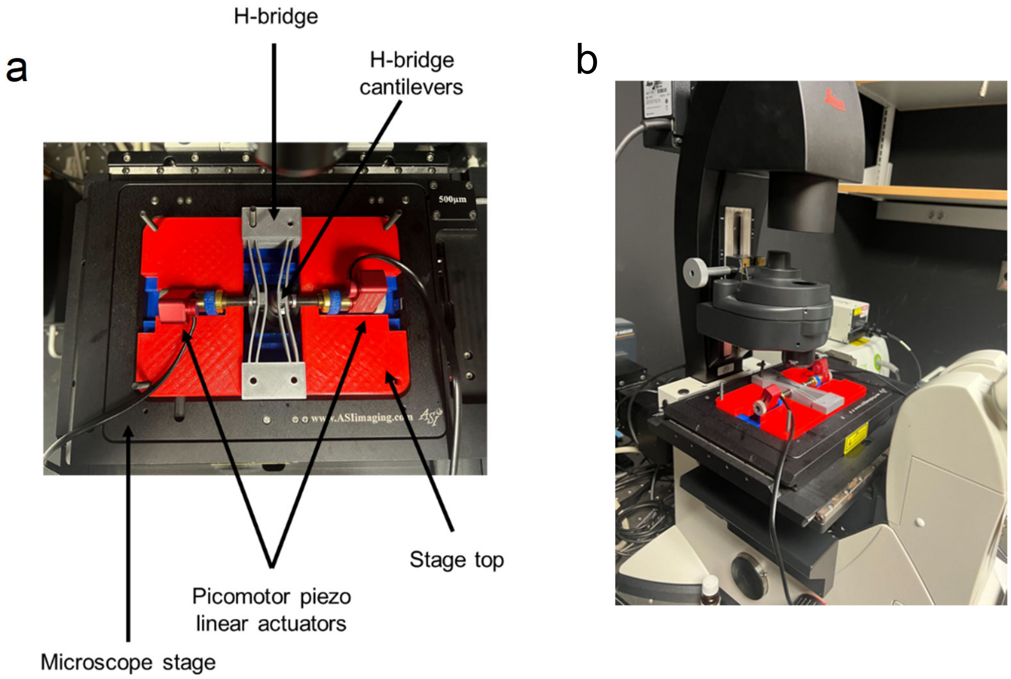


**3D-printed assembled microscope stage insert on a custom-built spinning disk confocal microscope.** (**a**) Top view of the stage insert on a microscope stage. The H-bridge sits on the stage top with crossbeams pushed inward by the picomotor piezo linear actuators that are mounted on the stage insert. (**b**) a diagonal view of the microscope stage insert.

**Supplementary Figure 2**

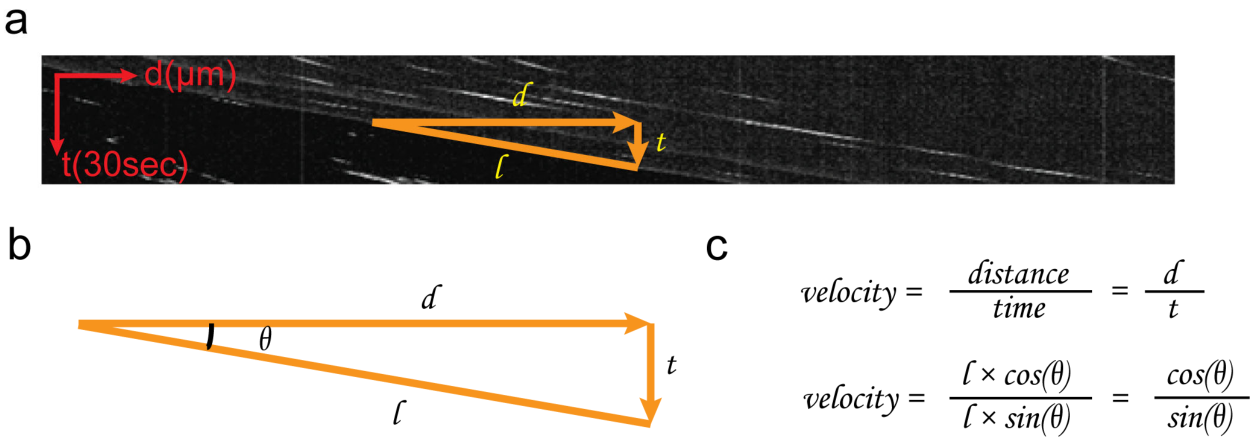


**Calculate the velocity of the edges of the cassette.** (**a**) Reslice of the timelapse video of one of the cassette edges, showing a kymograph-like trajectory of the edge traveled in distance (horizontal axis) and time (vertical axis). The edge appeared to travel in a straight line, indicating constant velocity during stretching. (**b**) A right triangle with an angle $\theta$ was used to calculate the velocity, with the horizontal leg as the distance $d$ and the vertical leg as the time $t$. The length of the hypotenuse was represented by $l$. (**c**) Velocity of the one side of the cassette edge was calculated by dividing distance by time. Velocity = 52 ± 3.5 μm/minute, n = 10.

**Supplementary Figure 3**

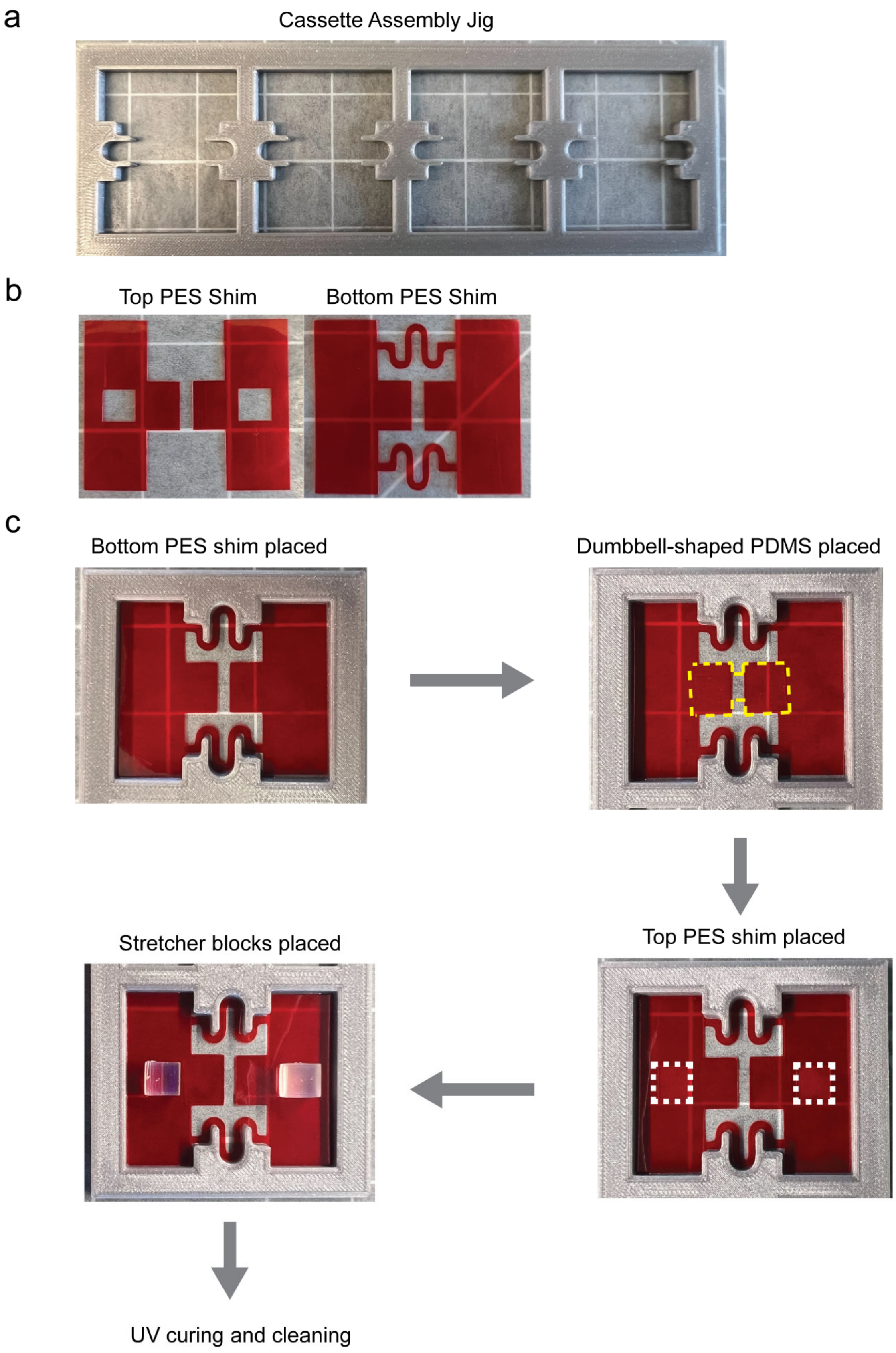

**Cassette Assembly using the 3D-printed jig.** (**a**) an FDM-printed jig with four cassette-shape compartments. (**b**) 2D-cut top PES shim with two cutouts for stretcher blocks alignment, and the bottom PES shim with spring cutouts. (**c**) Assemble the cassette using the jig. The bottom PES shim was placed in one of the compartments; a dumbbell-shaped PDMS sheet was placed onto the middle part of the cassette (yellow dashed line) with UV-curable optical adhesive; then the top PES shim was placed onto the cassette, followed by stretcher blocs with the optical adhesive (white dashed boxes indicated the cutouts for stretcher blocks). Then the assembled cassette was cured in UV chamber and washed with 100% ethanol and double deionized water.

**Supplementary Figure 4**

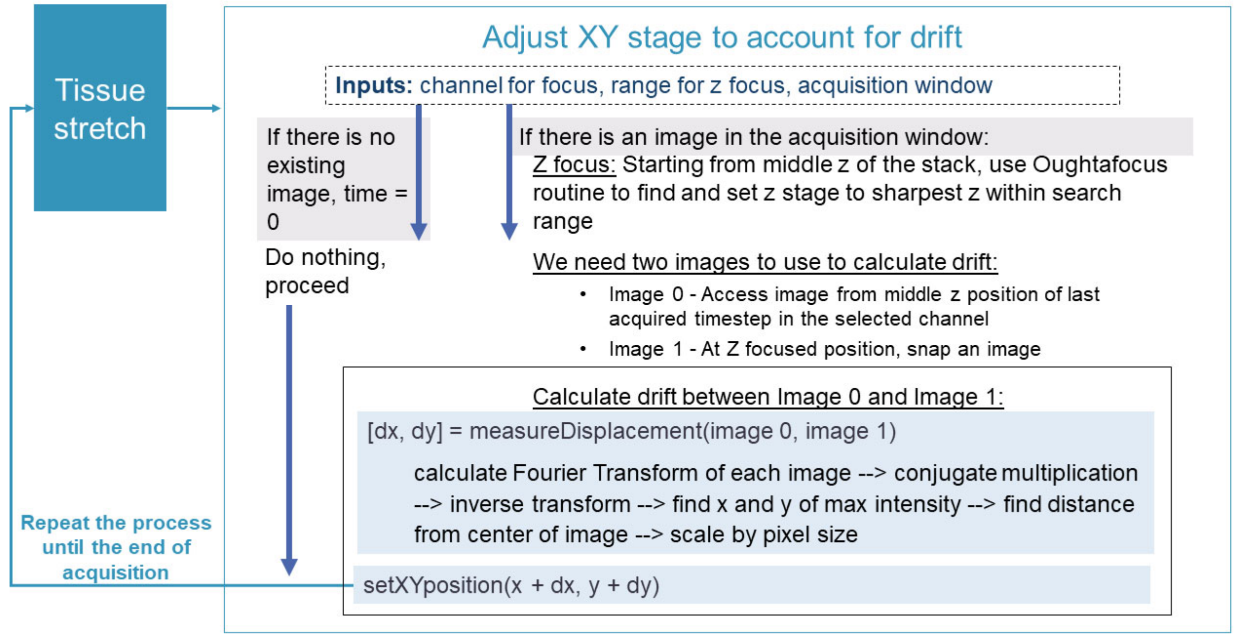


**A flowchart for customized AutoCenter plugin.** Between stretches of the motor, the xy stage is adjusted to account for drifting of the tissue that occurs during stretching. This is implemented with a custom autofocus plugin for Micro Manager, which handles image acquisition through ImageJ. This plugin uses the core autofocus functions, making it appear in the autofocus menu. The plugin module to adjust the xy stage to account for drift is executed at the beginning of each specified timepoint of image acquisition. The user supplies two inputs: the channel to focus in and the um range to use for z focus at the beginning of each timestep. First, the acquisition window is checked for an image. If no image exists, the plugin will do nothing and wait for the next round of images. If there is an image in the acquisition window, the Oughtafocus routine will be run from the middle of the stack plus and minus half the search range. This sharpest z is used to set the z position of the stage. Two images are needed to calculate the displacement. The first is taken from the middle z of the stack for the previous timepoint. The second is snapped at the newly focused z and stored within the plugin so it does not interfere with the acquisition. The displacement between these two images is calculated by conjugate multiplication of the Fourier Transforms of the images, inverse transforming the result, and then finding the deviation of the brightest pixel from the center of the image. This calculated dx and dy is then multiplied by the pixel size and used to set the new stage position.

**Supplementary Figure 5**

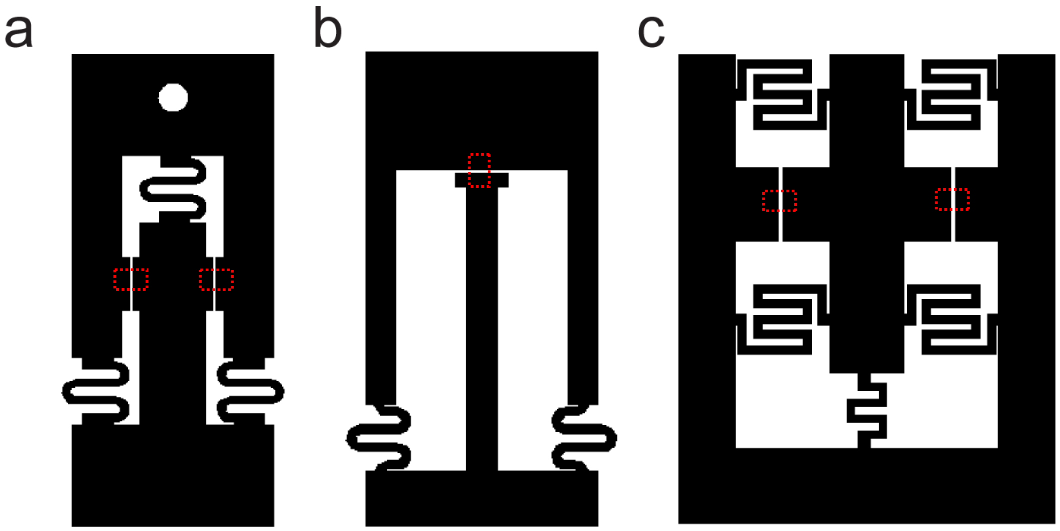


**Possible cassette designs for different strain profiles.** (**a**) a design for pure shear strain; (**b**) a design for unilateral stretch; (**c**) a design for biaxial stretch and shearing. Red boxes indicate substrates attachment.

**Supplementary Video 1**

Video of the top and bottom views of the stretcher system during stretching process. The cassette was in a relaxed state with the crossbeams of the H-bridge bent inward. As the arms of the linear actuators retracted, crossbeams straightened and stretched the cassette.

**Supplementary Video 2**

The cassette coated with green fluorescing polymer beads was stretched for 45 minutes. Images were taken every 30 seconds.

**Supplementary Video 3**

A fixed convallaria microscope slide was giving a simulated drift and imaged every 30 seconds for 7 minutes. **Left**: convallaria was shifted by the simulated drift without correction. **Right**: Same drift was applied but the AutoCenter plugin corrected the drift during acquisition.

**Supplementary Video 4**

Two stage 13 *Xenopus laevis* animal cap organotypic explants were stretched and imaged on an inverted brightfield microscope with a 2.5x objective. Images were taken every 18 seconds for 87 minutes.

**Supplementary Video 5**

A stage 13 *Xenopus laevis* animal cap organotypic explant expressed with membrane-mNeonGreen was stretched and imaged on an inverted spinning disk confocal microscope with a 25x water-immersion objective. Images were taken for 8 stretch steps (S0 – S8).

**Supplementary Video 6**

Mouse neonatal cardiomyocytes were imaged after stretch on an inverted spinning disk confocal microscope with a 63x oil-immersion objective. Cardiomyocytes remained intact and continued to beat after stretching. Images were taken every 15 seconds for 10 minutes.

Magenta: membrane; cyan: nuclei.
